# A choanoflagellate cGLR-STING pathway reveals evolutionary links between bacterial and animal immunity

**DOI:** 10.1101/2025.09.04.674280

**Authors:** Yao Li, Hunter C. Toyoda, Samantha G. Fernandez, Bianca Tiwari, Collin McNairy, Arielle Woznica, Philip J. Kranzusch

## Abstract

Animal innate immunity evolved from ancient pathways in bacterial anti-phage defense. How bacterial immune components were first acquired and adapted within eukaryotic cells remains poorly understood. Here we identify a complete cGLR-STING signaling axis in choanoflagellates, the closest living relatives of animals, that exhibits a mosaic of features from both bacterial and animal immunity. Comparative genomics reveals choanoflagellate *cGLR* and *STING* genes organized in operon-like arrangements reminiscent of bacterial defense loci. Reconstitution of choanoflagellate cGLR-STING signaling *in vitro* demonstrates that activation occurs through the conserved nucleotide immune signal 2′3′-cGAMP. Structural analysis of a choanoflagellate STING–2′3′-cGAMP complex explains how retention of bacterial-like features in early eukaryotic proteins shapes ligand specificity and receptor activation. We analyze *cGLR* and *STING* evolution in unicellular eukaryotes and identify further STING homologs in choanoflagellates and fungi that support additional independent acquisition events. Our results reveal molecular fossils that bridge bacterial and animal immunity and illuminate early eukaryotic immune system evolution.

## Introduction

The innate immune system of animals is comprised of many components that originated first in bacteria billions of years ago as ancient mechanisms of anti-phage defense^1-6^. As a founding example, the human cGAS-STING (cyclic GMP-AMP synthase, stimulator of interferon genes) pathway is an evolutionary descendent of bacterial defense operons named CBASS (cyclic oligonucleotide-based anti-phage signaling systems)^7-9^. In bacterial CBASS immunity, cGAS-like enzymes (named CD-NTases) detect phage infection and then synthesize nucleotide immune signals including 3′3′cGAMP (3′–5′ / 3′–5′ cyclic GMP-AMP) that bind to and activate downstream effector proteins that block phage propagation by inducing host cell death or growth ^8-13^. Some CBASS operons encode STING-like effector proteins that share direct structural and functional homology with human and animal STING^9,14-17^. In human cells, cGAS functions as a cytosolic DNA sensor that produces the related nucleotide immune signal 2′3′-cGAMP (2′–5′ / 3′–5′ cyclic GMP-AMP) and signals through STING-dependent activation of the type I interferon pathway^18-25^. Human cells therefore preserve the core signaling logic inherited from bacteria, while adapting it to regulate innate immunity and broadly conserved antiviral and antitumor responses.

In bacteria, CBASS immune components are organized into operons, with *CD-NTase* and *STING* genes positioned in close proximity to enable rapid activation of a linear signaling cascade in response to phage infection^8,9,26^. In contrast, metazoan cells often encode multiple distinct cGAS-like receptor (cGLR) and STING proteins that are no longer genomically linked and instead are distributed across distinct chromosomal loci (Fig. 1a)^27^. Individual animal genomes encode cGLRs with distinct immune specificities including proteins that recognize dsDNA, dsRNA, and yet unknown classes of ligands^27-29^. Additionally, they often encode multiple STING proteins that can exhibit distinct binding specificities to respond to endogenously synthesized cyclic dinucleotides including 2′3′-cGAMP and bacterial signals such as 3′3′-c-di-GMP and 3′3′-c-di-AMP^19,27^. Comparative genomic analyses across animal species have demonstrated a correlation in *cGLR* and *STING* gene distribution and abundance (Fig. 1b), supporting the hypothesis that both proteins were horizontally acquired from bacteria on the animal stem lineage^15,30^. A functional STING protein has been reported in the choanoflagellate species *Monosiga brevicollis*, one of the closest living relatives of animals, but the *in vivo* function of choanoflagellate cGLR proteins has remained elusive, and *in vitro* functional studies of putative cGLRs have not been performed for any unicellular eukaryote^31^. The lack of experimental models capable of providing insight into acquisition of cGLR and STING components into primitive eukaryotic genomes has limited understanding of early events in animal immune system evolution.

**Fig. 1.**
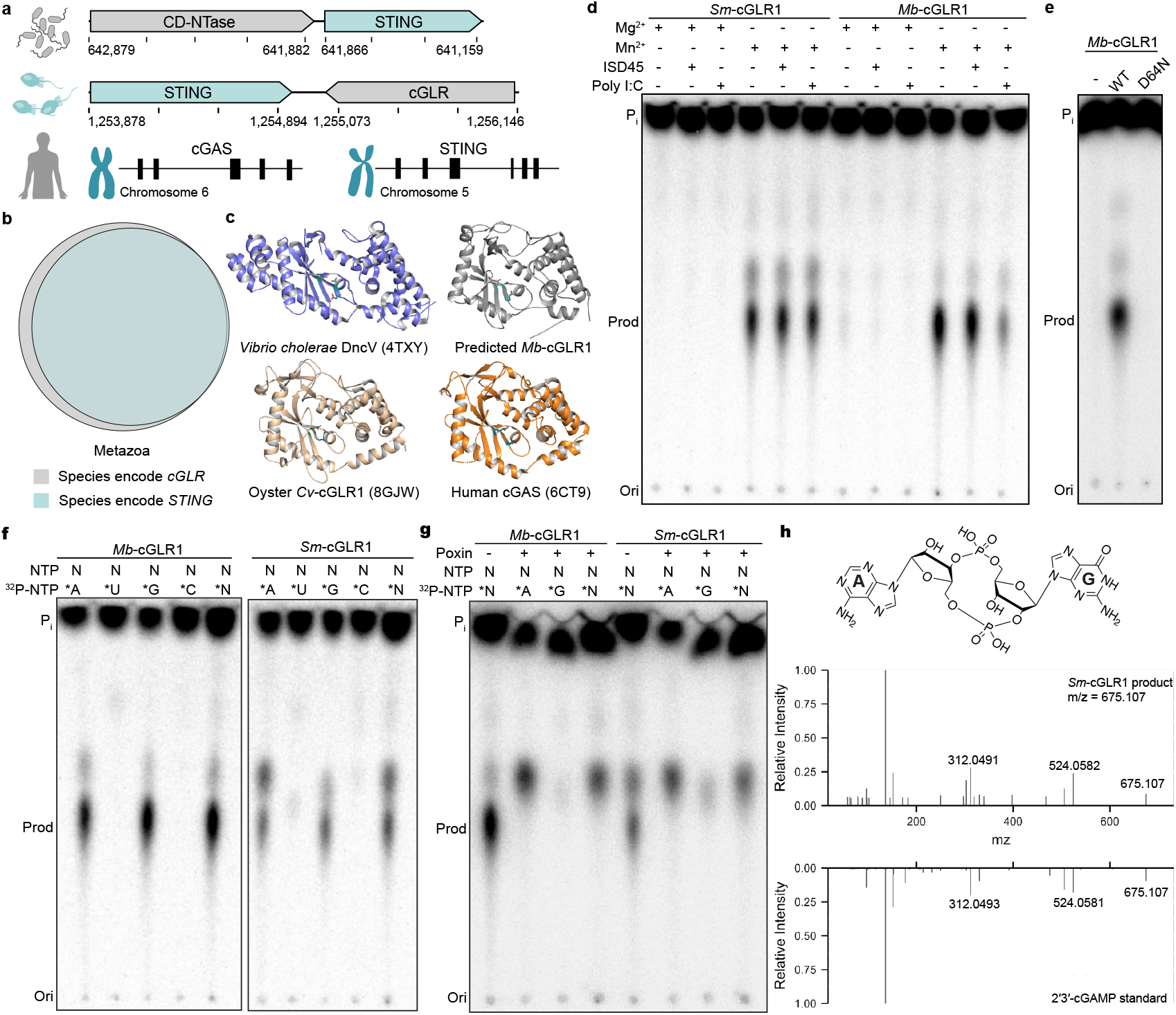
Discovery of functional cGLRs in Choanoflagellates. **a**, Configurations of *cGAS-like enzyme* and *STING* genes in the bacteria *S. faecium*, choanoflagellate *M. brevicollis*, and human genomes. In *S. faecium, CD-NTase* and *STING* genes are 4 basepairs apart from each other in the CBASS operon, which is responsible for defending against bacteriophage infection. In *M. brevicollis* genome, the *cGLR* and *STING* genes are colocalized with 178 basepairs in-between, while *cGAS* and *STING* genes are located on distinct chromosomes in human genome. **b**, Venn diagrams depicting the distribution of species encoding *cGLR* and *STING* genes in metazoans. Gene counts for each species are provided in Supplementary Table 1. **c**, Compared with previously published structures of the bacterial CD-NTase DncV (PDB: 4TXY)^39^, the oyster *C. virginica* cGLR (PDB: 8GJW)^27^, and human cGAS (PDB: 6CT9)^34^, the AlphaFold3-predicted structure of *Mb*-cGLR1 reveals an N-terminal NTase domain fused to a C-terminal five-helix bundle, resembling the overall architecture of cGAS-like enzymes. Catalytic residues (E62, D64, and D142 in *Mb*-cGLR1) are highlighted in teal. **d**, Thin layer chromatography of choanoflagellates *S. macro* and *M. brevicollis* cGLR activity in the presence of different divalent metal ions (magnesium or manganese) and/or ligands (a 45 bp dsDNA or polyI:C dsRNA). Both cGLRs are insensitive to double-stranded DNA or RNA. Data are representative of n = 3 independent experiments. **e**, The catalytic activity of *Mb-*cGLR1 is abolished when one of the key residues in the [E/D]h[E/D] X_50–90_ [E/D] catalytic triad, D64, is mutated to asparagine. Data are representative of n = 3 independent experiments. **f**, Thin layer chromatography analysis of *Mb-*cGLR1 and *Sm-*cGLR1 reactions labeled with individual α^32^P-NTPs. CIP (calf intestinal phosphatase) was used to terminate the reactions by removing terminal phosphate groups from nucleotides. The results suggest both cGLRs utilize GTP and ATP to produce a nucleotide second messenger. Data are representative of n = 3 independent experiments. **g**, Thin layer chromatography analysis of reactions from panel **f** further treated with poxin, a phosphodiesterase that specifically cleaves the 3′–5′ phosphodiester bond in 2′3′-cGAMP. The results indicate both choanoflagellate cGLRs likely produces 2′3′-cGAMP. Data are representative of n = 3 independent experiments. **h**, MS/MS analysis of *Sm-* cGLR1 product in comparison to synthetic 2′3′-cGAMP standard.

Here, we define a complete cGLR-STING signaling axis in choanoflagellates that reveals a remarkable blend of evolutionary features derived from both bacterial- and metazoan-like immune proteins. We identify multiple choanoflagellate species that encode *cGLR* and *STING* genes in a single syntenic arrangement that mirrors ancient operon organization in bacterial CBASS immunity. Biochemical analyses demonstrate that choanoflagellate cGLRs specifically synthesize the nucleotide immune signal 2′3′- cGAMP and that the associated choanoflagellate STING proteins exhibit correspondingly selective recognition of 2′3′- cGAMP as an activating signal. By combining enzymatic, structural, and phylogenetic analyses, we uncover how retention of bacterial-like features in early eukaryotic STING proteins shapes ligand specificity and the mechanism of Li *et al*. 2025 (preprint) 2 receptor activation. We build on these findings to identify additional clades of standalone STING proteins in choanoflagellates, fungi, and other unicellular eukaryotes and expand analysis of early evolution of cGLR-STING pathways into eukaryotic cells. Our findings illuminate how prokaryotic immune modules were reconfigured during early eukaryotic evolution and establish microbial eukaryotes as a powerful system for probing the origins of animal immune signaling.

## Results

### Discovery of a functional cGLR in choanoflagellates

To provide insight into the evolution of animal innate immunity, we searched early-branching eukaryotic genomes for potential cGLR-STING signaling pathways. In bacteria, primitive STING proteins function in anti-phage defense and are encoded adjacent to CD-NTase enzymes that control cyclic dinucleotide synthesis and upstream signal initiation^9,15^. We therefore hypothesized that some eukaryotic genomes may retain this linked genetic architecture and that *STING* gene neighborhoods may be enriched for related immune signaling enzymes. Previously, Woznica *et al*. identified a functional STING protein in the single cell choanoflagellate species *M. brevicollis* (*Mb*STING) that is necessary for mediating responses to the nucleotide signal 2′3′-cGAMP^31^. We analyzed the *M. brevicollis* genome and identified immediately downstream of *MbSTING* a candidate nucleotidyltransferase (NTase)-superfamily signaling enzyme encoded by the uncharacterized gene *MONBRDRAFT_36418* (Fig. 1a). Although *MONBRDRAFT_36418* runs in an antisense head-to-tail orientation with *Mb*STING (Fig. 1a), and previous transcriptome analysis indicated these genes are expressed at different levels^31,32^, AlphaFold3^33^ predictions of the *MONBRDRAFT_36418*-encoded protein supported conservation of the bi-lobed architecture characteristic of animal cGLR and bacterial CD-NTase immune signaling proteins and the presence of a complete [E/D]h[E/D] X50–90 [E/D] required for synthesis of a cyclic dinucleotide immune signal (Fig. 1c)^7,27,28,34^. We identified homologous proteins in the transcriptomes of four other choanoflagellate species (Extended Data Fig 1a)^32^, including a protein with 32% aminoacid identity encoded adjacent to *STING* in *S. macrocollata* (*S. macro, Sm*STING), demonstrating that this nucleotidyl-transferase protein occurs in multiple genomes as a STING-associated gene (Fig. 1a, Extended Data Fig 1b).

To investigate the function of STING-associated nucleotidyltransferase proteins, we expressed and purified the choanoflagellate *M. brevicollis* and *S. macro* gene products and tested the recombinant proteins for enzymatic activity *in vitro* (Extended Data Fig.1c–e). Using radiolabeled nucleotide substrates and thin-layer chromatography, we observed that both nucleotidyltransferase proteins were active enzymes capable of synthesizing a nucleotide product and we therefore renamed these proteins *M. brevicollis* cGLR1 (*Mb*-cGLR1) and *S. macro* cGLR1 (*Sm*-cGLR1) (Fig. 1d). Human cGAS and many animal cGLRs function as nucleic acid sensors that initiate enzymatic activity when in complex with stimulating DNA and RNA ligands^18,27,28,35^. In contrast, *Mb*-cGLR1 and *Sm*-cGLR1 activity does not require the presence of DNA or RNA (Fig. 1d), suggesting that these enzymes may function more similar to bacterial CD-NTase enzymes that are also constitutively active *in vitro*^7^. The activities of *Mb*-cGLR1 and *Sm*-cGLR1 each required the presence of the divalent cation Mn^2+^ and were abolished by mutations to the conserved cGLR active-site triad required for metal coordination (Fig. 1d–e).

We next combined biochemical and mass spectrometry profiling to identify the *Mb*-cGLR1 and *Sm*-cGLR1 product nucleotide signal. Using reactions supplemented with individual α^32^P-radiolabeled NTPs, we observed that the major product of both choanoflagellate cGLRs was specifically labeled with GTP and ATP (Fig. 1f). The final product was resistant to phosphatase treatment consistent with synthesis of a cyclic dinucleotide signal (Fig. 1f). We labeled the individual guanine and adenine nucleobases and treated the products with Nuclease P1, a nuclease that promiscuously cleaves 3′−5′ linkages but is unable to cleave 2′−5′ linkages, and observed that the adenosine phosphate is incorporated into a protected 2′−5′ bond consistent with synthesis of the nucleotide signal 2′−5′ / 3′−5′ cyclic GMP-AMP (2′3′-cGAMP) (Extended Data Fig 1f). To confirm these findings, we treated the *Mb*-cGLR1 and *Sm*-cGLR1 products with poxin, an animal viral immune evasion nuclease that specifically recognizes 2′3′-cGAMP and cleaves the 3′−5′ bond, and observed complete degradation (Fig. 1g). Finally, tandem mass spectrometry profiling (MS/MS) compared to a synthetic standard confirmed that the *Mb*-cGLR1 and *Sm*- cGLR1 product nucleotide signal is 2′3′-cGAMP (Fig. 1h, Extended Data Fig 1g–j). Together, these results demonstrate that choanoflagellate species encode a functional cGLR signaling enzyme that produces the nucleotide signal 2′3′- cGAMP.

### Choanoflagellate *Sm*STING specifically recognizes 2′3′- cGAMP

Animal cGLRs signal through activation of the downstream receptor STING^18,27,28^. We therefore tested the ability of the choanoflagellate cGLR1 product 2′3′-cGAMP to interact with choanoflagellate STING. We expressed and purified the cyclic dinucleotide binding domain of *Sm*STING (Extended Data Fig 2a–b) and observed using an electrophoretic mobility shift assay that *Sm*STING forms a stable complex with 2′3′-cGAMP and that *Sm*STING–2′3′-cGAMP complex formation occurs with low micromolar binding affinity of ∼2.5 µM (Fig. 2a–b). In bacterial anti-phage defense, STING cyclic dinucleotide receptors exhibit highly specific binding and typically only recognize the specific nucleotide immune signal produced by an associated CD-NTase enzyme^9,16,17^. In animal innate immunity, STING cyclic dinucleotide receptors typically recognize both endogenous cGLR nucleotide immune signals like 2′3′-cGAMP and 2′3′-cUA as well as prokaryotic cyclic dinucleotides including 3′3′-c- di-GMP and 3′3′-c-di-AMP^21,24,27,36^. To define the ligand specificity of choanoflagellate STING we analyzed the ability of *Sm*STING to interact with a panel of radiolabeled 2′3′- and 3′3′-linked cyclic dinucleotides. Unlike most metazoan STING receptors, *Sm*STING is highly selective for 2′3′- cGAMP and does not interact with other cyclic dinucleotides including the closely related regioisomer 3′2′-cGAMP (Fig. 2b–c, Extended Data Fig 2c). These results explain previous *in vivo* observations that choanoflagellate *M. brevicollis* cells respond to 2′3′-cGAMP stimulation, but not bacterial cyclic dinucleotides 3′3′-c-di-GMP or 3′3′-di-AMP^31^. This suggests that the early branching eukaryotic cGLR-STING pathway in choanoflagellates exhibits characteristics similar to CD-NTase-STING proteins in bacterial anti-phage defense.

**Fig. 2.**
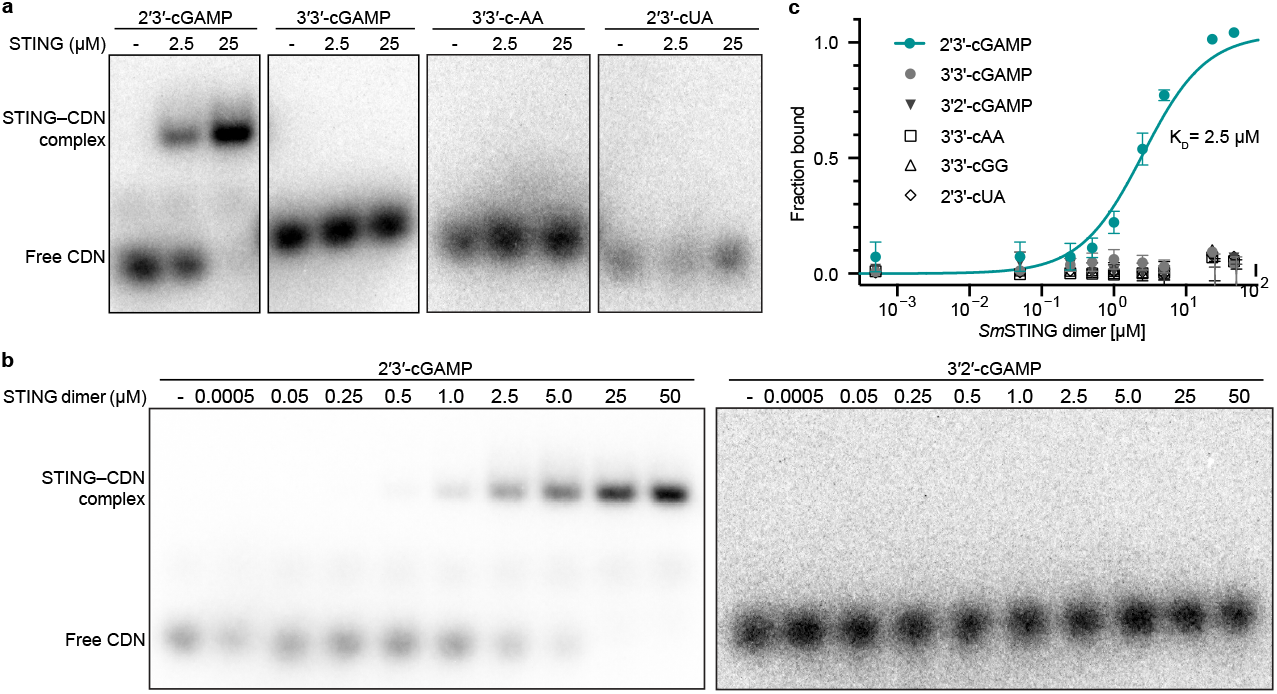
Choanoflagellate *Sm*STING specifically recognizes 2′3′-cGAMP. **a**, Electrophoretic mobility shift assay (EMSA) analysis of the binding of *Sm*STING to 2′3′-cGAMP, 3′3′-cGAMP, 3′3′-cAA and 2′3′-cUA performed at STING dimer concentrations of 0, 2.5, and 25 µM. Data are representative of n = 2 independent experiments. Information on the purification of *Sm*STING are included in Extended Data Fig 2a–b. **b**, Primary EMSA analysis of two isomers of cGAMP, 2′3′-cGAMP and 3′2′-cGAMP, titrated with increasing amount of *Sm*STING protein spanning a concentration ranging from 0.5 nM to 50µM (homodimer form). Primary EMSA analysis of titration of other CDNs with *Sm*STING are included in Extended Data Fig 2c. Data are representative of n = 2 independent experiments. **c**, Quantification of the EMSA analysis in panel **b** and Extended Data Fig 2c. Only 2′3′-cGAMP demonstrated detectable binding to *Sm*STING. The binding affinity was determined by nonlinear regression analysis of the binding curve. Data are the mean ± std of n = 2 independent experiments.

### Structure of choanoflagellate SmSTING reveals features of bacterial and mammalian immunity

Choanoflagellate STING proteins have a predicted architecture similar to human STING with four N-terminal transmembrane helices fused to a C-terminal cyclic dinucleotide-binding domain (Extended Data Fig 3a). The C-terminal cyclic dinucleotide-binding domains of *Sm*STING and *Mb*STING are 29% identical at the amino acid level, supporting significant divergence over evolutionary time in these Craspedid choanoflagellate species^37^. To gain insight into the evolution of choanoflagellate STING and define the molecular determinants of 2′3′-cGAMP selectivity, we purified *Sm*STING fused to a stabilizing T4 lysozyme domain and determined a 2.7 Å crystal structure of the protein in complex with 2′3′-cGAMP (Fig. 3a, Extended Data Fig 3b–d and Supplementary Table 2).

**Fig. 3.**
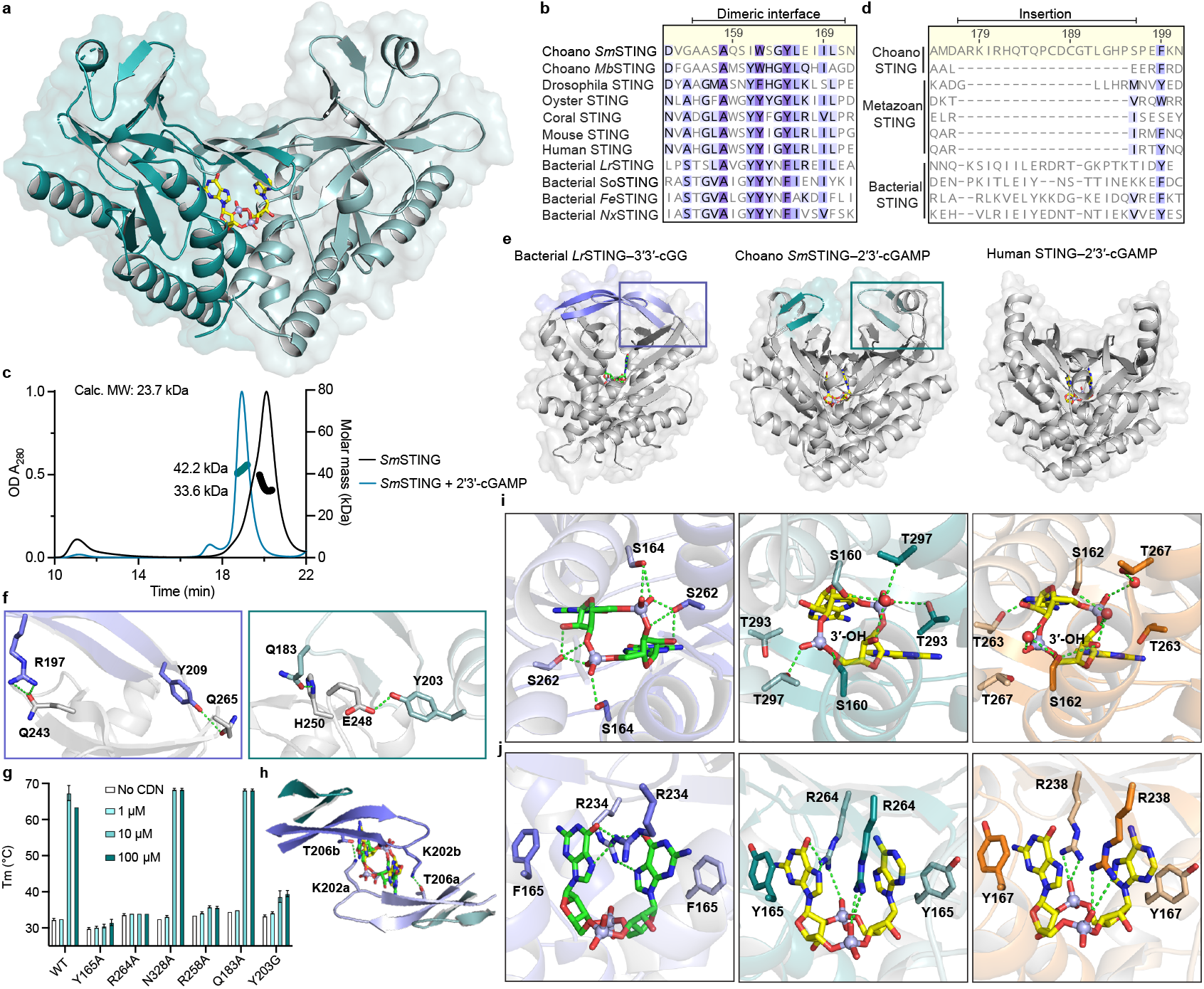
Structure of choanoflagellate SmSTING reveals features of bacterial and mammalian immunity. **a**, Crystal structure of *Sm*STING C-terminal cyclic dinucleotide binding domain in complex with 2′3′-cGAMP. **b**, Sequence alignment of choanoflagellate STING receptors with STING proteins from representative metazoan and bacterial species reveals a conserved hydrophobic patch that facilitates dimerization. **c**, Size-exclusion chromatography coupled with multi-angle light scattering (SEC-MALS) revealed that *Sm*STING ΔTM undergoes a monomer-to-dimer transition upon binding 2′3′-cGAMP. **d** and **e**, Sequence alignment and structural comparison of bacterial (PDB 8HWI)^17^, choanoflagellate and human STING (PDB 4KSY)^24^ receptors highlighting an additional β-strand turn (β1–β2) in *Sm*STING, a bacterial STING-like structural motif absents from all characterized metazoan STING homologs. **f**, Cutaway view showing hydrogen bonds between the β-strand turn extension and canonical lid region in *Lr*STING and *Sm*STING. Colored boxes correspond to the framed region in panel **e. g**, Thermofluor assays measuring the thermal stability (*T*_m_) of six *Sm*STING mutants compared to wild type (WT) in the presence of increasing concentrations of 2′3′-cGAMP (0, 1, 10, 100 µM) (n = 3). An upward shift in *T*_m_ indicates ligand-induced stabilization, while reduced or absent shifts relative to WT reflect impaired 2′3′-cGAMP recognition (see Methods for assay details). **h**, Top view of *Lr*STING illustrating β-strand turn-mediated inter-protomer hydrogen bonds between T206 and K202 that stabilize the long-lid conformation. These interactions are absent in *Sm*STING due to the shorter β-turn insertion. **I** and **j**, Structural comparison of bacterial *S. faecium* STING–3′3′-c-di-GMP (purple, PDB 7UN8)^14^, *Sm*STING–2′3′-cGAMP (teal), and human STING–2′3′-cGAMP (orange, PDB 4KSY)^24^ highlighting conserved hydrogen-bonding networks and bacterial-like nucleobase-specific interactions in *Sm*STING.

The structure of choanoflagellate STING reveals a surprising combination of features previously thought to be unique to bacterial or metazoan STING proteins. The *Sm*STING cyclic dinucleotide-binding domain forms a canonical V-shaped, homodimeric architecture that is conserved in all STING proteins across the tree of life (Fig. 3a). At the base of the homodimeric interface, *Sm*STING helix α1 forms a hydrophobic stem known to be required for dimerization of human and metazoan STING proteins (Fig. 3b, Extended Data Fig 3e)^21,24,27,36^. However, the *Sm*STING stem sequence ASAQSIW is distinct from the canonical GLAWSYY motif in human and metazoan STING proteins and exhibits increased hydrophilic character more similar to bacterial STING stem helices (Fig. 3b, Extended Data Fig 3e). In agreement with a potentially weakened dimeric interface, in the absence of ligand the *Sm*STING cyclic dinucleotide-binding domain predominantly exists as a monomer in solution distinct from the obligate dimers observed with human and metazoan STING proteins (Fig. 3c, Extended Data Fig 3f)^21,24,27,36^. The *Sm*STING ΔTM protein dimerizes only in the presence of 2′3′- cGAMP, not with other cyclic dinucleotides (Extended Data Fig 3g–j), consistent with the ligand binding affinity analysis (Fig. 2c). A STING monomer-to-dimer transition was previously reported for several bacterial STING cyclic dinucleotide-binding domains, including those from *Roseivirga ehrenbergii* and *Flavobacteriaceae sp*., suggesting that similar to bacterial receptors *Sm*STING may require ligand-induced dimerization for activation^9^. The *Sm*STING structure also reveals an additional beta strand turn β1–β2 previously observed in some bacterial STING proteins that extends the canonical β-strand lid region responsible for sealing the ligand-binding pocket (Fig. 3d–e)^17^. Notably, this bacterial STING-like structural motif is absent from all characterized metazoan STING homologs^21,24,27,36^. In both *Sm*STING and bacterial *Larkinella arboricola* STING (*Lr*STING), the additional β-strand turn extension forms two pairs of hydrogen bonds with residues on the underlying canonical lid (Fig. 3f). We introduced a Y203G mutation to disrupt this hydrogen bonding interaction in *Sm*STING and observed reduced 2′3′-cGAMP binding affinity, supporting a conserved role for this motif in stabilizing the protein-ligand complex (Fig. 3g, Extended Data Fig 4a–c). In some bacterial STING proteins like *Lr*STING, the extended β- strand lid forms inter-protomer hydrogen bonds between K202 and T206, which have been shown to contribute to autoinhibition and reduce cytotoxicity^17^. In *Sm*STING, the β- strand lid is shorter and cannot form equivalent inter-protomer interactions (Fig. 3h), suggesting that choanoflagellates may employ alternative modes of STING regulation.

To further explore how bacterial STING-like characteristics may influence choanoflagellate STING ligand recognition, we next analyzed *Sm*STING interactions with 2′3′-cGAMP. In bacterial STING proteins, ligand selectivity is often driven by nucleobase-specific interactions, whereas in human and other metazoan STING proteins recognition is primarily mediated through hydrogen bonds with the cyclic dinucleotide’s phosphodiester backbone, leading to a strong preference for mixed 2′–5′/3′–5′-linked backbones^21,24^. A key distinction lies in the role of a conserved arginine residue: in bacterial STING (R234 in *Sf*STING), this residue recognizes the guanine bases of 3′3′-c-di-GMP, while in human and metazoan STING proteins (R238 in human STING), this residue is repurposed to coordinate the phosphate backbone^9,21,24^. In the *Sm*STING ligand binding pocket, residues S160, T262, and T266 form a hydrogen-bonding network with 2′3′-cGAMP that is remarkably similar to interactions in the binding pocket of human STING (Fig. 3i). However, the conserved arginine residue R264 instead makes nucleobase-specific contacts to the guanosine base in 2′3′-cGAMP, mirroring the nucleobase-specific interactions observed in bacterial STING (Fig. 3j, Extended Data Fig 4d)^9,14^. Mutagenesis analysis of the *Sm*STING cyclic dinucleotide-binding pocket confirmed the critical role of R264 and other conserved residues in controlling 2′3′- cGAMP recognition (Fig. 3g, Extended Data Fig.4e–i). Together, these structural and biochemical analyses demonstrate that choanoflagellate STING proteins share an overall architecture conserved with animal STING proteins and reveal how retention of bacterial STING-like features contributes to selective ligand recognition.

### Evolution of STING function in early-branching eukaryotes

We next used the structure of *Sm*STING and analysis of choanoflagellate proteins to examine the evolution of STING from bacterial to metazoan immunity. We identified 50 *STING* genes in diverse early-branching eukaryotic genome and transcriptome sequences that retained all structural features of the *Sm*STING cyclic dinucleotide binding domain required for ligand recognition (Supplementary data 1). Notably, analysis of these sequences revealed additional choanoflagellate STING proteins encoded in genomes that lack a detectable *cGLR* gene suggesting a function for STING outside of the canonical cGLR-STING signaling axis (Extended Data Fig 1a). To evaluate the function of choanoflagellate STING proteins that occur in the absence of cGLR signaling, we expressed and purified two example STING proteins from the choanoflagellates *Stephanoeca diplocostata* and *Diaphanoeca grandis* and tested the ability of these proteins to bind a panel of cyclic dinucleotide ligands. Surprisingly, *Stephanoeca diplocostata* and *Diaphanoeca grandis* STING did not bind 2′3′-cGAMP and instead exhibited specific recognition of the cyclic dinucleotide 3′3′-c-di-GMP, a widely conserved nucleotide second messenger molecule in bacteria (Fig. 4a)^38^. Selective recognition of 3′3′-c-di-GMP suggests that these choanoflagellate receptors retain ancestral ligand specificity inherited from bacterial STING proteins that function in 3′3′-c-di-GMP-dependent anti-phage defense systems^9^. Both *Stephanoeca diplocostata* and *Diaphanoeca grandis* STING proteins bound 3′3′-c-di-GMP with a dissociation constant of approximately 1 µM (Fig. 4b, Extended Data Fig 4j), consistent with values reported for bacterial STING proteins.

**Fig. 4.**
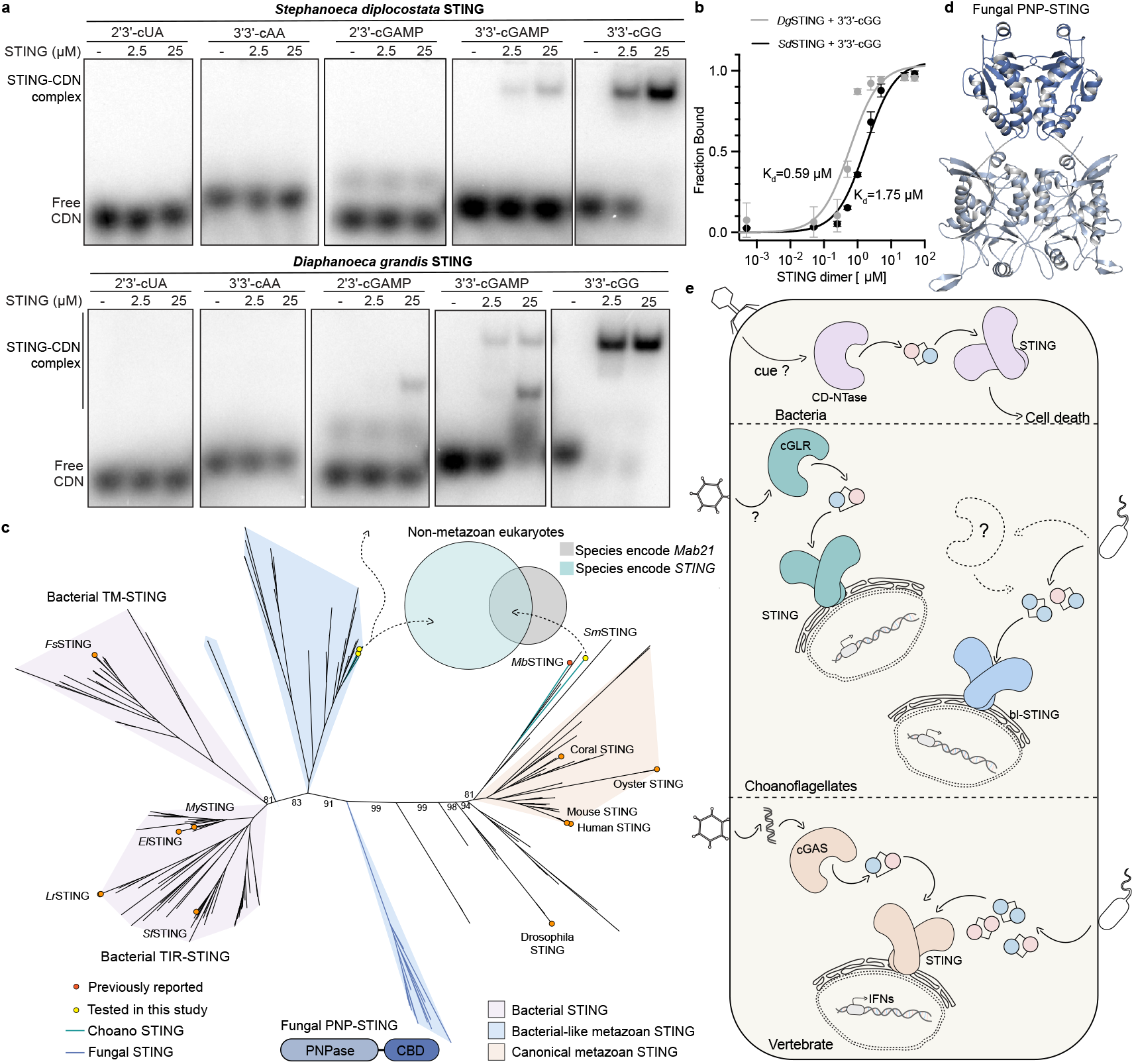
Evolution of STING function in early-branching eukaryotes. **a**, EMSA analysis of bacterial-like STING receptors from choanoflagellates *S. diplocostata* and *D. grandis* binding to 2′3′-cUA, 3′3′-cAA, 2′3′-cGAMP, 3′3′-cGAMP, and 3′3′-c-di-GMP, performed at STING dimer concentrations of 0, 2.5, and 25 µM. Data are representative of n = 2 independent experiments. **b**, Quantification of the EMSA analysis in Extended Data Fig 4j showing specific recognition of 3′3′-c-di-GMP with dissociation constants (*K*_d_) of 0.59 µM for *Dg*STING and 1.75 µM for *Sd*STING. **c**, Maximum-likelihood phylogenetic tree of representative bacterial and eukaryotic STING proteins, including choanoflagellate, stramenopile, amoebozoan, fungal, and metazoan sequences. Choanoflagellate STING proteins recognizing 3′3′-c-di-GMP cluster near bacterial STING homologs, whereas cGLR-associated choanoflagellate STING proteins (e.g., *Sm*STING, *Mb*STING) group with metazoan homologs. A distinct clade of fungal PNP-STING proteins forms an isolated branch that groups more closely with bacterial STING receptors. Bootstrap values were calculated in RAxML^40^ and are shown at the nodes. The Venn diagram shows the distribution of species encoding STING and Mab21 superfamily genes in non-metazoan eukaryotes. Gene counts for each species are from Cumberson and Levin (2023)^30^. Details of the bioinformatic analysis are provided in the Methods. **d**, AlphaFold3-predicted structure of a representative fungal PNP-STING (from *Zasmidium cellare*) showing the canonical STING homodimer fold, conserved dimeric interface, and lid-region insertions reminiscent of bacterial STING receptors. The STING cyclic dinucleotide binding domain (CBD) is shown in a dark blue, and the fused N-terminal PNPase domain is shown in a lighter shade. **e**, Model for the evolution of STING signaling in eukaryotes, showing an inferred founding event in which a complete cGLR–STING signaling axis was incorporated into animal innate immunity, alongside independent acquisitions of bacterial-like STING modules for 3′3′-c-di-GMP sensing in other eukaryotic lineages.

To explain these alternative roles for STING in eukaryotic cells, we built upon previous structure-guided analyses of bacterial and metazoan STING cyclic dinucleotide binding domain sequences and constructed a maximum-likelihood phylogenetic tree of STING homologs from representative prokaryotic and eukaryotic lineages^9,30,31^. The resulting phylogeny revealed STING proteins from early-branching eukaryotes including choanoflagellate, stramenopile, amoebozoa, and fungi lineages form a continuum of STING branches that bridge the evolutionary space between bacterial and metazoan STING proteins (Fig.4c). Interestingly, choanoflagellate STING proteins that recognize 3′3′-c-di-GMP and those that function as part of 2′3′-cGAMP-dependent cGLR-STING signaling axes form divergent branches on the phylogenetic tree (Fig. 4c). Choanoflagellate STING proteins that recognize 3′3′-c-di-GMP cluster near bacterial homologs, consistent with a previously described group of bacterial-like eukaryotic STING proteins^30^, while those partnered with *cGLR* genes, such as *Sm*STING and *Mb*STING, align more closely with metazoan STING proteins consistent with a signaling arrangement similar to animal innate immunity (Fig. 4c).

Surprisingly, our analysis also uncovered a previously unrecognized clade of fungal STING homologs that forms a distinct and isolated branch in the phylogenetic tree positioned well apart from all previously characterized bacterial and metazoan STING proteins (Fig. 4c). While these fungal sequences do not cluster closely with any known STING group, they branch closer to bacterial STING proteins than to metazoan homologs, suggesting a possible ancestral relationship with prokaryotic signaling modules. In bacteria, STING domains are often fused to effector modules such as an enzymatic TIR (Toll/interleukin-1 receptor) domain, coupling cyclic dinucleotide binding to antiviral defense through cellular growth arrest^9^. In contrast, the fungal STING homologs identified here are not associated with transmembrane or TIR domains, but instead are fused to a purine nucleoside phosphorylase (PNP) domain, a fusion not previously reported in characterized STING proteins. An AlphaFold3^33^-predicted model of a representative fungal PNP-STING protein supports a canonical STING homodimeric fold and shared cyclic dinucleotide binding domain (Fig. 4d). The absence of detectable *cGLR* genes in fungal genomes suggests that PNP-STING proteins may also operate independently of canonical cGLR-STING signaling pathways. Given the deep phylogenetic placement, structural similarity to bacterial homologs, and unique domain architecture, these fungal STING proteins may have resulted from an independent acquisition of prokaryotic signaling components into eukaryotes, consistent with the modular reformulation of prokaryotic immune systems in diverse eukaryotic lineages^15^. Although limited by the sparse coverage of currently sequenced eukaryotic species, the separate distribution of eukaryotic STING proteins suggests a model where potential independent evolutionary events give rise to standalone sensors of microbial nucleotide signals like 3′3′- c-di-GMP and a rarer founding event of acquisition of a complete anti-phage defense-like operon that led to adaptation of cGLR-STING signaling axes in animal immunity (Fig. 4e). These findings explain early emergence of STING in primitive unicellular eukaryotes and provide insight into the evolutionary flexibility of STING signaling architectures in eukaryotic immunity.

## Discussion

Our results define a functional cGLR-STING signaling axis in choanoflagellates and demonstrate that unicellular eukaryotes can preserve both the biochemical logic and genomic organization of immune systems originating in bacterial antiphage defense (Fig. 4e). Using biochemical and structural analyses, we discover that multiple choanoflagellate species encode cGLRs that synthesize 2′3′- cGAMP and that adjacent STING receptors selectively recognize 2′3′-cGAMP as an activating ligand (Fig.s. 1 and 2). Although independently regulated in choanoflagellates, the arrangement of *cGLR* and *STING* genes in choanoflagellates mirrors the operon-like organization of CBASS loci in bacteria and provides further insight into the evolutionary link between bacterial anti-phage defense and animal innate immunity (Fig. 1a)^7-9^.

Analysis of choanoflagellate cGLR and STING proteins reveals a mosaic of features from both bacterial and metazoan immune signaling. Choanoflagellate cGLR proteins exhibit constitutive activity *in vitro* similar to many bacterial CD-NTase enzymes in CBASS anti-phage defense (Fig. 1d)^7^. However, choanoflagellate cGLR proteins function more similarly to metazoan cGLR immune proteins and selectively synthesize the nucleotide immune signal 2′3′-cGAMP common throughout animal immunity^27^. Choanoflagellate cGLR activity is likely controlled in cells to induce immune signaling but future *in vivo* studies in choanoflagellates will be required to further define the mechanism of pathway activation. We find that choanoflagellate STING proteins use a similar mixture of bacterial- and metazoan-like features to control ligand recognition. The choanoflagellate STING cyclic dinucleotide-binding pocket recognizes 2′3′-cGAMP with a hydrogen-bonding pattern shared with human and animal STING proteins that coordinates high-affinity recognition of the ligand mixed 2′–5′ / 3′–5′ phosphodiester backbone (Fig. 3i)^21,24^. Additional bacterial STING-like interactions in choanoflagellate STING form further arginine-nucleobase contacts that make the binding site highly specific for 2′3′- cGAMP and unable to recognize other cyclic dinucleotides commonly observed as ligands in many animal STING proteins (Fig. 2c)^19,21,27^. The choanoflagellate STING cyclic dinucleotide binding pocket also retains bacterial STING-like extensions in the β-strand lid region suggesting that this element is retained for structural stability or may serve an alternative regulatory function preserved in unicellular eukaryotic cells. Together, these data support a model in which choanoflagellate cGLR and STING proteins exhibit intermediate features retained during the transition of proteins from bacterial anti-phage defense to metazoan innate immunity.

Choanoflagellate cGLR-STING proteins additionally provide insight into proposed models of early evolution of animal immunity. In both *M. brevicollis* and *S. macrocollata* choanoflagellates, the *cGLR* and *STING* genes are positioned adjacent in the genome, consistent with bacterial-like synteny and supporting a shared horizontal acquisition from prokaryotes. We additionally identified *STING* genes present in choanoflagellate genomes that lack detectable cGLR proteins (Extended Data Fig 1a). Analysis of cyclic dinucleotide binding specificity demonstrates that these STING proteins do not recognize 2′3′-cGAMP and instead retain bacterial STING-like high-affinity recognition of 3′3′- c-di-GMP (Fig. 4a–b). 3′3′-c-di-GMP is a common signal in nearly all bacterial phyla, suggesting that acquisition of standalone *STING* genes may have enabled some choanoflagellate species to detect foreign bacterial ligands. Finally, our structure-guided analysis of unicellular STING proteins also identified a clade of fungal STING proteins that occur as fusions with PNP effector-like domains (Fig. 4c–d). These fungal STING proteins also occur in genomes lacking detectable cGLR signaling enzymes, and branch closer to bacterial STING proteins than to metazoan homologs, suggesting further independent acquisition events from bacteria.

Together, our results support a unifying model in which multiple, independent horizontal gene transfer events introduced STING and other early immune proteins into eukaryotic genomes. Following these rare events in evolution, subsequent modifications result in currently observable present-day features in eukaryotes including lineage-specific fusions, regulatory ligand adaptations, and integration into broader signaling networks. The cGLR-STING pairs in choanoflagellates provide a possible example of remnants of ancient intact acquisitions where immune systems with complete structural and functional continuity are retained from the bacterial anti-phage defense origins. Standalone genes, like cGLR-independent STING homologs in choanoflagellates and PNP-STING proteins in fungi, likely represent separate acquisition events in eukaryotes and illustrate how immune pathways are repeatedly captured and modified to meet specific cellular functions. Further exploration of immune proteins in microbial eukaryotes will provide a path to uncover unexpected functional innovations and explain the evolutionary processes that have shaped immune signaling networks across the tree of life.

## Acknowledgements

The authors are grateful to members of the Kranzusch laboratory for helpful comments and discussion. We thank the Center for Macromolecular Interactions at Harvard Medical School and the Dana-Farber Cancer Institute Molecular Biology Core Facility. X-ray data were collected at beamline 17-ID-1 of the National Synchrotron Light Source II (NSLS2), a US Department of Energy (DOE) Office of Science User Facility operated under Contract No. DE-SC0012704. X-ray data were additionally collected with support from an agreement between the Advanced Photon Source (a U.S. DOE Office of Science User Facility operated by Argonne National Laboratory under Contract No. DE-AC02-06CH11357) and the Diamond Light Source, the UK’s national synchrotron facility located at the Harwell Science and Innovation Campus in Oxfordshire. The work was funded by grants to P.J.K. from the Pew Biomedical Scholars program, the Burroughs Wellcome Fund PATH program, The G. Harold and Leila Y. Mathers Charitable Foundation, The Mark Foundation for Cancer Research, the Cancer Research Institute, the Parker Institute for Cancer Immunotherapy, and the National Institutes of Health (1DP2GM146250-01) and a grant to A.W. from the Howard Hughes Medical Institute (Hanna Gray Faculty Fellows Program). Y.L. is supported by the Parker Institute for Cancer Immunotherapy.

## Author contributions

The study was designed and conceived by Y.L. and P.J.K. Y.L. performed all biochemistry and nucleotide signal analysis experiments with assistance from H.C.T. and B.T. Structural biology experiments were performed by H.C.T. and Y.L. S.G.F., C.M. and A.W. analyzed *cGLR* and *STING* genomic operons sequences. cGLR and STING protein sequence bioinformatic analysis was performed by Y.L. The manuscript was written by Y.L. and P.J.K. All authors contributed to editing the manuscript and support the conclusions.

## Data Availability Statement

All data are available in the manuscript or the supplementary information. The crystal structure of SmSTING has been deposited in the Protein Data Bank under accession code 9Q1F.

## Additional Information

Correspondence and requests for materials should be addressed to P.J.K. All illustrations were created using Adobe Illustrator.

## Materials and Methods

### Cloning and plasmid construction

*Mb*-*cGLR1, Sm*-*cGLR1* and *SmSTING ΔTM* genes were synthesized as gBlocks (Integrated DNA Technologies) with ≥18 base pairs of homology flanking the insert sequence and cloned into a custom pETSUMO2 or pET-SUMO2-T4lysosome vector by Gibson assembly as previously described^28,34^. Plasmids of *Sm*STING mutants were generated from the parental plasmid pETSUMO2-*Sm*STING using the one-step site-directed plasmid mutagenesis protocol developed by Liu *et al*^41^. Briefly, each mutagenic primer comprised a 5′ primer–primer overlapping region (∼10–15 nt) carrying the desired substitution and a longer 3′ non-overlapping region to promote primer–template annealing. The Tm of the non-overlap region was set ≥10°C higher than the Tm of the overlap region. PCR reactions (25 µl) were assembled with 5 ng plasmid template, 1.25 µl of each primer (10 µM), 5 µl Q5 Reaction Buffer, 5 µl Q5 GC Enhancer, 2.5 µl dNTPs (2 mM each), and 0.25 µl Q5 DNA polymerase (New England Biolabs). The annealing temperature was set according to the Tm of the non-overlapping region of the primers. Amplified products were digested with DpnI (37°C, 2 h) to remove methylated parental plasmid and transformed into *E. coli* Top10. All mutations were confirmed by Sanger sequencing.

### Protein expression and purification

cGLR and STING proteins were expressed as recombinant proteins as previously described^27,34^. Briefly, expression plasmids were transformed into *E. coli* strain BL21-RIL (Agilent) and grown overnight in 30 mL MDG media at 37°C with shaking at 230 RPM. Overnight MDG cultures were used to inoculate 2–4 L M9ZB medium, which was grown at 37°C until OD_600_ reached ∼2.5. Protein expression was then induced with 0.5 mM IPTG, and cultures were incubated overnight at 16°C with shaking at 230 rpm. Bacterial cells were harvested by centrifugation, resuspended and lysed in lysis buffer (20 mM HEPES-KOH pH 7.5, 400 mM NaCl, 30 mM imidazole, 10% glycerol and 1 mM DTT). Lysates were cleared by centrifugation and loaded onto Ni-NTA resin (Qiagen), washed with lysis buffer supplemented with 1 M NaCl and eluted with lysis buffer supplemented with 300 mM imidazole. The SUMO2 tag was removed by incubating with recombinant human SENP2 protease during dialysis into a buffer containing 20 mM HEPES-KOH pH 7.5, 250 mM KCl, 1 mM DTT and 10% glycerol. *Sm*STING mutants were concentrated to >10 mg mL^−1^, flash frozen with liquid nitrogen and stored in −80°C. *Mb*-cGLR1 and *Sm*-cGLR1 proteins were further purified with by ion-exchange using a 5 mL HiTrap Heparin HP column (Cytiva) and eluted with a NaCl gradient from 150 mM to 1 M. The resulting cGLR protein fractions, together with *Sm*STING ΔTM and T4-lysosome– *Sm*STING ΔTM from Ni-NTA elution, were subsequently purified by size-exclusion chromatography on a 16/600 Superdex 75 or 16/600 Superdex 200 column (Cytiva). Final protein preparations were concentrated to >10 mg mL^−1^, flash frozen with liquid nitrogen, and stored at −80°C.

### *In vitro* cGLR second messenger synthesis and nucleotide purification

The nucleotide synthesis activity of *Mb*-cGLR1 and *Sm*-cGLR1 was assessed by thin-layer chromatography (TLC) as described previously^27,28^. Briefly, 1 µL purified cGLR protein was incubated at 37°C overnight in reaction buffer (50 mM Tris-HCl pH 7.5, 100 mM KCl, 10 mM MgCl_2_ or 1 mM MnCl_2_, 1 mM DTT) containing 50 µM each unlabeled ATP, CTP, GTP, and UTP, and 0.5 µL of α-^32^P-labeled NTPs (∼0.4 µCi each). Reactions were performed in the absence of ligand or in the presence of 1 µg poly I:C or 5 µM ISD45 dsDNA as indicated. Reactions were terminated by adding 0.5 µL Quick CIP (NEB) to remove terminal phosphates from unreacted nucleotides. Aliquots (0.5 µL) were spotted on 20 × 20 cm PEI-cellulose TLC plates (Millipore), developed in 1.5 M KH_2_PO_4_ pH 3.8 until the solvent front was 1–3 cm from the top, air-dried, and imaged on a Typhoon Trio Variable Mode Imager (GE Healthcare) after exposure to a phosphor screen.

To determine the nucleotide composition of cGLR products, 0.25–1 µM cGLR protein was incubated with 50 µM each unlabeled NTP and 0.5 µL of a single α-^32^P-labeled NTP. Reactions were terminated with Quick CIP and analyzed by TLC as above. For poxin degradation assays, 4 µL reaction was incubated with 1 µL of 10 µM poxin at 37°C for 1 h. For nuclease P1 assays, 4 µL reaction was treated with 0.5 µL nuclease P1 (Sigma N8630) and 0.5 µL Quick CIP in 1× P1 buffer (30 mM NaOAc pH 5.3, 5 mM ZnSO_4_, 50 mM NaCl) for 1 h at 37°C, then analyzed by TLC.

To collect cGLR nucleotide products for mass spectroscopy analysis, 100 µL reactions were prepared in the same conditions as described above. The reaction mixtures were treated with Quick CIP for 3 h and heated for 1 h at 65°C before centrifugation at 4°C, 3,200 × g for 10 min and filtration through a 0.22 µm filter to remove precipitated protein. Samples were then spun through a 10-kDa molecular weight cut off spin column (Amicon) to remove protein and high molecular weight ligand. HPLC analysis was carried out as previously described^27,28^ at 40°C using a C18 column (Agilent Zorbax Bonus-RP 4.6×150 mm, 3.5-micron) with a mobile phase of 50 mM NaH_2_PO_4_ (pH 6.8 with NaOH) supplemented with 3% acetonitrile and run at 1 mL/min. Nucleotide products of cGLRs were collected based on retention time using the fraction collector of the HPLC instrument (Agilent 1200 series) and concentrated using a speed vac before mass spectrometry analysis.

### Liquid chromatography-tandem mass spectrometry (LC-MS/MS) analysis

LC-MS/MS analysis samples were analyzed by the commercial company MS-Omics (now part of Clinical Microbiomics) as previously described^27,28^. Analysis was carried out using a Vanquish™ Horizon UHPLC System coupled to Orbitrap Exploris 240 Mass Spectrometer (Thermo Fisher Scientific, US). The UHPLC was performed using an Infinity Lab PoroShell 120 HILIC-Z PEEK lined column with the dimension of 2.1 x 150mm and particle size of 2.7µm (Agilent Technologies). The composition of Mobile phase A was 10 mM ammonium acetate at pH 9 in 90% Acetonitrile LC-MS grade (VWR Chemicals, Leuven) and 10 % Ultra-pure water from Direct-Q® 3 UV Water Purification System with LC-Pak® Polisher (Merck KGaA, Darmstadt) and mobile phase B was 10 mM ammonium acetate at pH 9 in ultra-pure water with 5 µM medronic acid (InfinityLab Deactivator additive, Agilent Technologies). The flow rate kept at 250 µl mL^−1^ consisting of a 2 min hold at 10% B, increased to 40% B at 14 min, held till 15 min, decreased to 10% B at 16 min and held for 8 min. The column temperature was set at 30°C and an injection volume of 5 µl.

A heated electrospray ionization interface was used as ionization source on the MS. Analysis was performed in positive and negative ionization mode from m/z 300 to 1500 at a mass resolution of 120000. Ion source parameters used: Sheath gas flow rate, 20 (arbitrary units); auxiliary gas flow rate, 5 (arbitrary units); Sweep gas flow rate, 1 (arbitrary units), capillary temperature, 350°C; S-lens radiofrequency level 70; automatic gain control (AGC) target, 1E6 (Standard); maximum injection time, 100 ms; spray voltage 2.5 kV in negative and 3.5 kV in positive. MS2 spectra was acquired using data dependent acquisition (DDA) with a mass resolution of 45000, an isolation window m/z 0.4 and normalized collision energy of 20, 40 and 60 eV. Data were manually inspected to generate MS/MS spectra using Freestyle 1.4 (Thermo Fisher Scientific).

### Crystallization and structure determination

Crystal of the *Sm*STING CBD–2′3′-cGAMP complex was grown at 18°C for 3–30 days using hanging-drop vapor diffusion. Purified *Sm*STING CBD fused to N-terminal T4 lysosome was diluted to 5 mg mL^−1^ in buffer (20 mM HEPES-KOH 7.5, 75 mM KCl, 1 mM TCEP) and incubated with 0.5 mM 2′3′-cGAMP on ice for 10 min. Initial screens were set up in 96-well trays (70 µL reservoir) by mixing 200 nL protein–ligand complex with 200 nL reservoir solution using a Mosquito liquid handler (SPT Labtech). Optimized crystals were obtained in EasyXtal 15-well trays (NeXtal Biotechnologies) with 400 µL reservoir solution, combining 1 µL protein solution with 1 µL reservoir solution. The optimized condition contained 5% ethylene glycol, 100 mM MOPS pH 6.65, and 10% (w/v) PEG8000. Crystals were cryoprotected with reservoir solution supplemented with 10– 25% ethylene glycol and harvested with a nylon loop.

X-ray diffraction data were collected at the NSLS2, beamline AMX, and processed using the autoPROC toolbox^42^. Experimental phases were determined by single-wavelength anomalous dispersion (SAD) from data collected on selenomethionine-substituted protein prepared as described previously^43^. Anomalous sites were identified, and an initial map was generated with AutoSol within PHENIX^44^. Structural modelling was completed in Coot^45^ and refined with PHENIX.

### Electrophoretic mobility shift assay

Electrophoretic mobility shift assays were performed to assess the interactions between STING proteins and cyclic dinucleotides as previously described^9,27^. Briefly, 20 nM of each α-^32^P labeled cyclic dinucleotide was incubated with STING proteins at the indicated concentrations or across a serial dilution range (0.5 nM–50 µM) in buffer containing 5 mM magnesium acetate, 50 mM Tris-HCl pH 7.5, 50 mM KCl, and 1 mM DTT. After incubation at room temperature for 10 min, reactions were resolved on a 7.2 cm, 6% native polyacrylamide gel run at 100 V for 45 min in 0.5× TB buffer. Gel was fixed in a solution of 40% ethanol and 10% acetic acid for 15 min before drying at 80°C for 40 min. The dried gel was exposed to a phosphorscreen and imaged on a Typhoon Trio Variable Mode Imager (GE Healthcare). Signal intensity was quantified using ImageQuant 5.2 software. The binding affinity was determined by nonlinear regression analysis of the binding curve, generated by plotting protein concentration (0–50 µM) against the percentage of bound ligand.

### Thermofluor assay

Thermal stability of *Sm*STING proteins was assessed by differential scanning fluorimetry (Thermofluor). Purified wild-type (WT) and mutant *Sm*STING ΔTM proteins (Y165A, R264A, N328A, R258A, Q183A, Y203G) were diluted to 15 µM in a buffer containing 20 mM Tris-HCl pH 7.5, 100 mM KCl and incubated with 2′3′-cGAMP at final concentrations of 0, 1, 10, or 100 µM in a total reaction volume of 20 µL. SYPRO Orange dye (4×; Invitrogen) was added to monitor protein unfolding. Samples were heated from 20°C to 95°C in a BioRad CFX thermocycler with HEX channel fluorescence measurements every 0.5°C. Melting temperatures (Tm) were determined from derivative plots (–dF/dT) of the fluorescence–temperature curves, with Tm defined as the temperature at which the derivative reached its minimum value. The resulting Tm values were plotted as bar graphs in GraphPad Prism.

### Bioinformatics and tree construction

The Venn diagram in Fig. 1b and Fig. 4c summarizing eukaryotic species encoding cGLR (or Mab21 superfamily) and STING genes was generated based on previously reported datasets^27,30^. Gene counts for each species are provided in Supplementary Table 1.

Initial searches of choanoflagellate STING genes were performed by manually running BLAST searches with E-value cutoff of 0.005 of the *Mb*STING cyclic dinucleotide-binding domain against published choanoflagellate transcriptomes^31^. Broader searches for STING homologs were carried out by constructing hidden Markov models (HMMs) from representative bacterial (*Fs*STING) and choanoflagellate (*Sm*STING) sequences, followed by iterative PSI-BLAST searches against the NCBI non-redundant (nr) protein database and BLAST searches against the EukProt v3 database^46^. Four PSI-BLAST iterations were run with an E-value cutoff of 0.005, BLOSUM62 scoring matrix, gap existence cost of 11, gap extension cost of 1, and conditional compositional score matrix adjustment enabled. BLAST searches against EukProt were performed with an E-value cutoff of 0.005^47^. Newly identified sequences were combined with sequences previously identified in Li *et al*. (2023)^27^ and Culbertson and Levin (2023)^30^ and truncated to the CBD region for comparative analysis. Multiple sequence alignments were generated with MUSCLE using the ppp algorithm in Geneious Prime v2025.2.1. Maximum-likelihood phylogenetic trees were constructed with RAxML-NG v1.2.2^40^ using the Le and Gascuel amino acid replacement model with a gamma distribution of rate variation among sites (LG+G) ^48^ and 500 bootstrap replicates. Trees were visualized and annotated in iTOL v6^49^ and taxonomic classifications were obtained from the metadata of each NCBI non-redundant protein entry. TMHMM 2.0^50^ was used to predict transmembrane segments.

### Statistics and reproducibility

Experimental details regarding replicates and sample size are described in the figure legends.

---

**Extended Data Figure 1.**
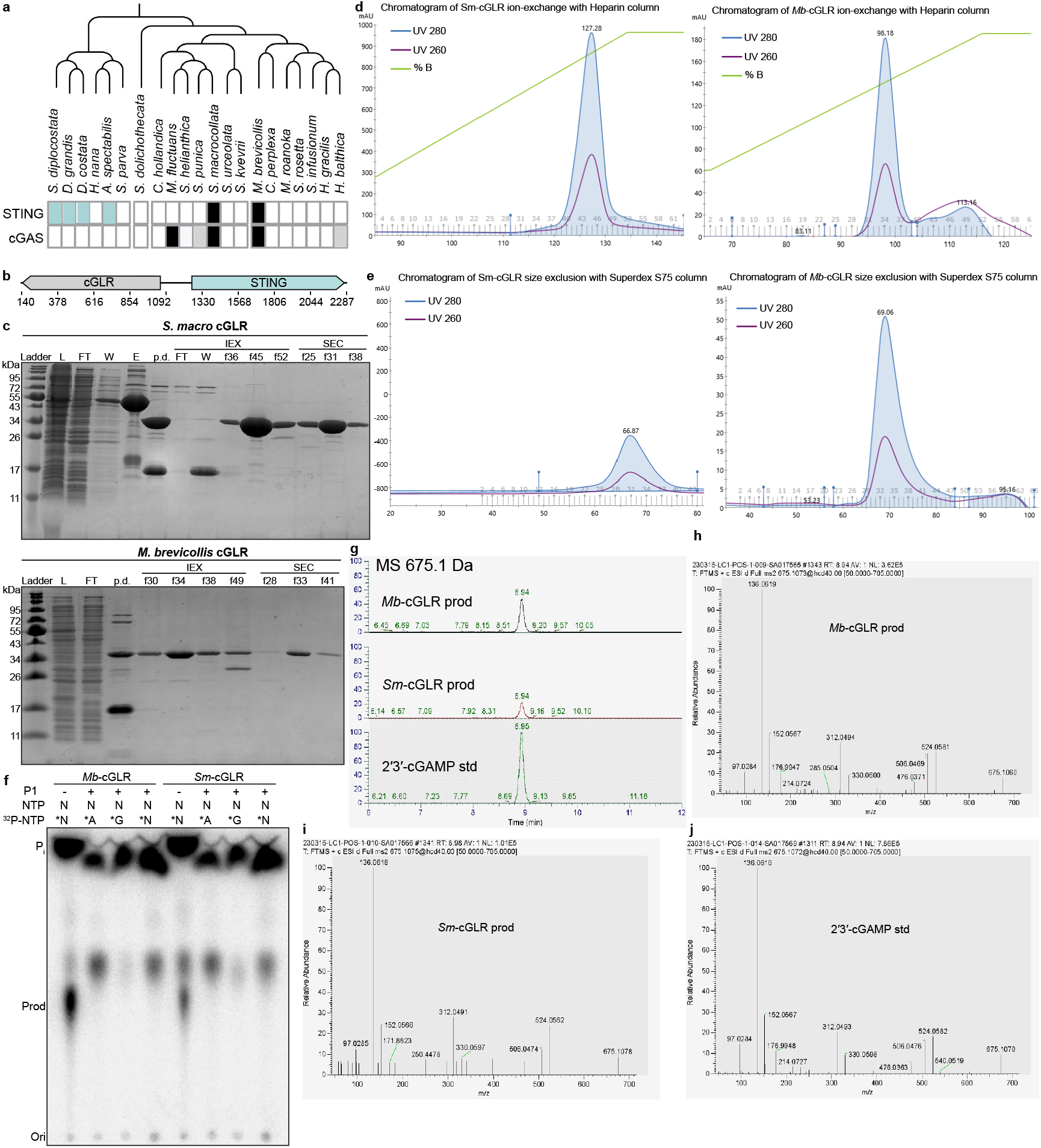
Genomic context and characterization of choanoflagellate cGLRs. **a**, Distribution of cGLR and STING genes across 21 choanoflagellate species, re-illustrated from Woznica *et al*. (eLife 2020)^31^ with newly discovered choanoflagellate cGLR and STING genes indicated in grey and light teal, respectively. Black boxes denote genes described in the original eLife study. **b**, Configurations of *cGLR* and *STING* genes in the choanoflagellate *Salpingoeca macrocollata*. **c–e**, Protein expression levels and purification of *Sm*-cGLR1 and *Mb*-cGLR1. Sodium dodecyl sulfate–polyacrylamide gel electrophoresis (SDS–PAGE) with Coomassie stain shows protein samples before and after purification steps (**c**), including Ni–NTA affinity chromatography, ion-exchange chromatography (**d**), and size-exclusion chromatography (**e**). **f**, Thin layer chromatography analysis of reactions from Fig. 1f further treated with nuclease P1, a phosphodiesterase that cleaves the 3′–5′ but not 2′–5′ phosphodiester bond in oligonucleotides. **g**, LC-MS analysis showing extracted ion chromatograms for the 675.1 Da [M+H]_+_ ion corresponding to 2′3′-cGAMP in *Mb-*cGLR1 and *Sm-*cGLR1 reaction products, compared to a synthetic 2′3′-cGAMP standard. **h–j**, LC-MS/MS fragmentation spectra of the *Mb*-cGLR1 product (**h**), *Sm*-cGLR1 product (**i**), and synthetic 2′3′-cGAMP standard (**j**), confirming identical fragmentation patterns consistent with 2′3′-cGAMP.

**Extended Data Figure 2.**
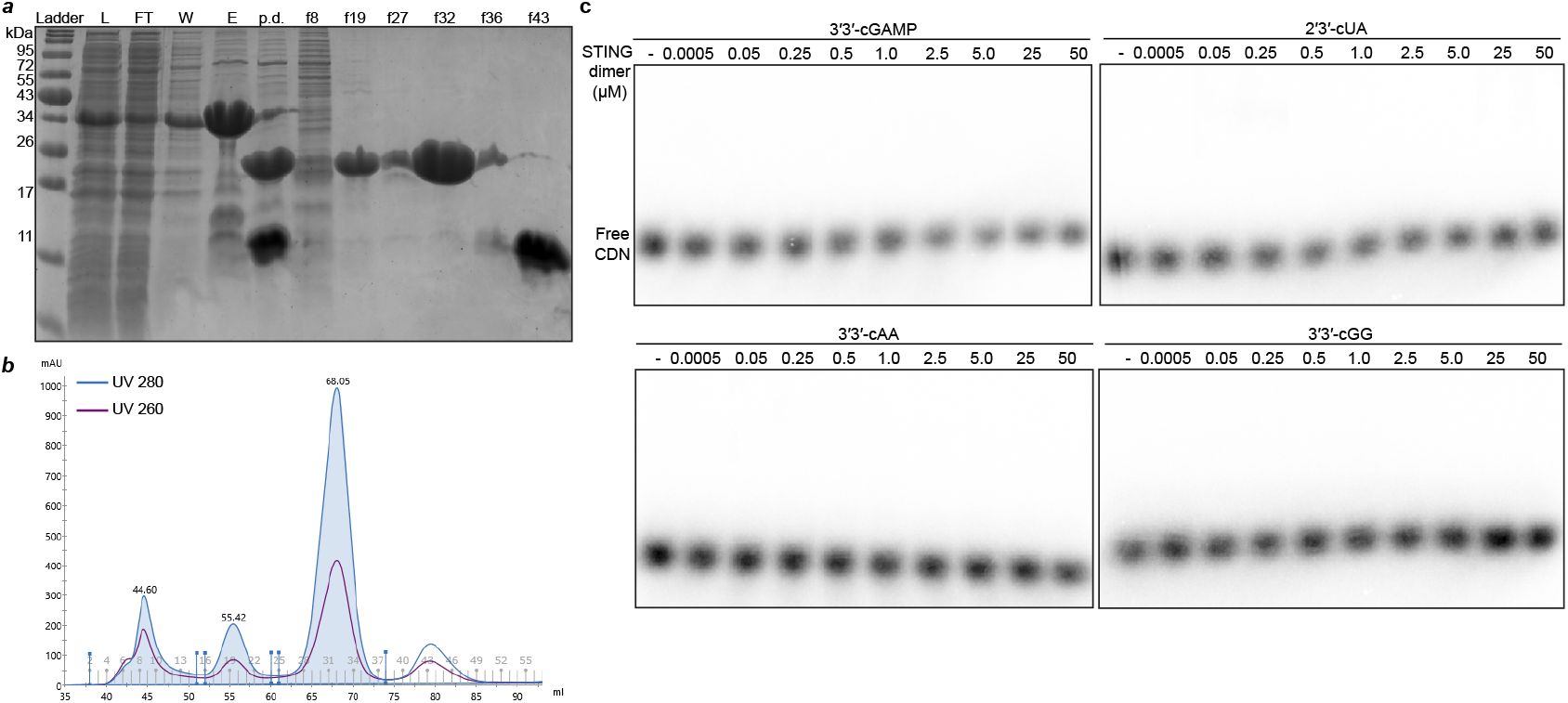
Electrophoretic mobility shift assay of *Sm*STING–ligand interactions. **a**–**b**, Purification of the cyclic dinucleotide-binding domain of *Sm*STING. SDS–PAGE with Coomassie stain (**a**) shows protein samples before and after Ni–NTA affinity chromatography and size-exclusion chromatography, with the corresponding size-exclusion chromatogram shown in (**b**). **c**, Primary EMSA analysis of other CDNs tested in this study, including 3′3′-cGAMP and 2′3′-cUA, 3′3′-cAA and 3′3′-cGG. Ligand binding was assessed by titrating *Sm*STING protein (homodimer form) over concentrations ranging from 0.5 nM to 50 µM. Data are representative of n = 2 independent experiments.

**Extended Data Figure 3.**
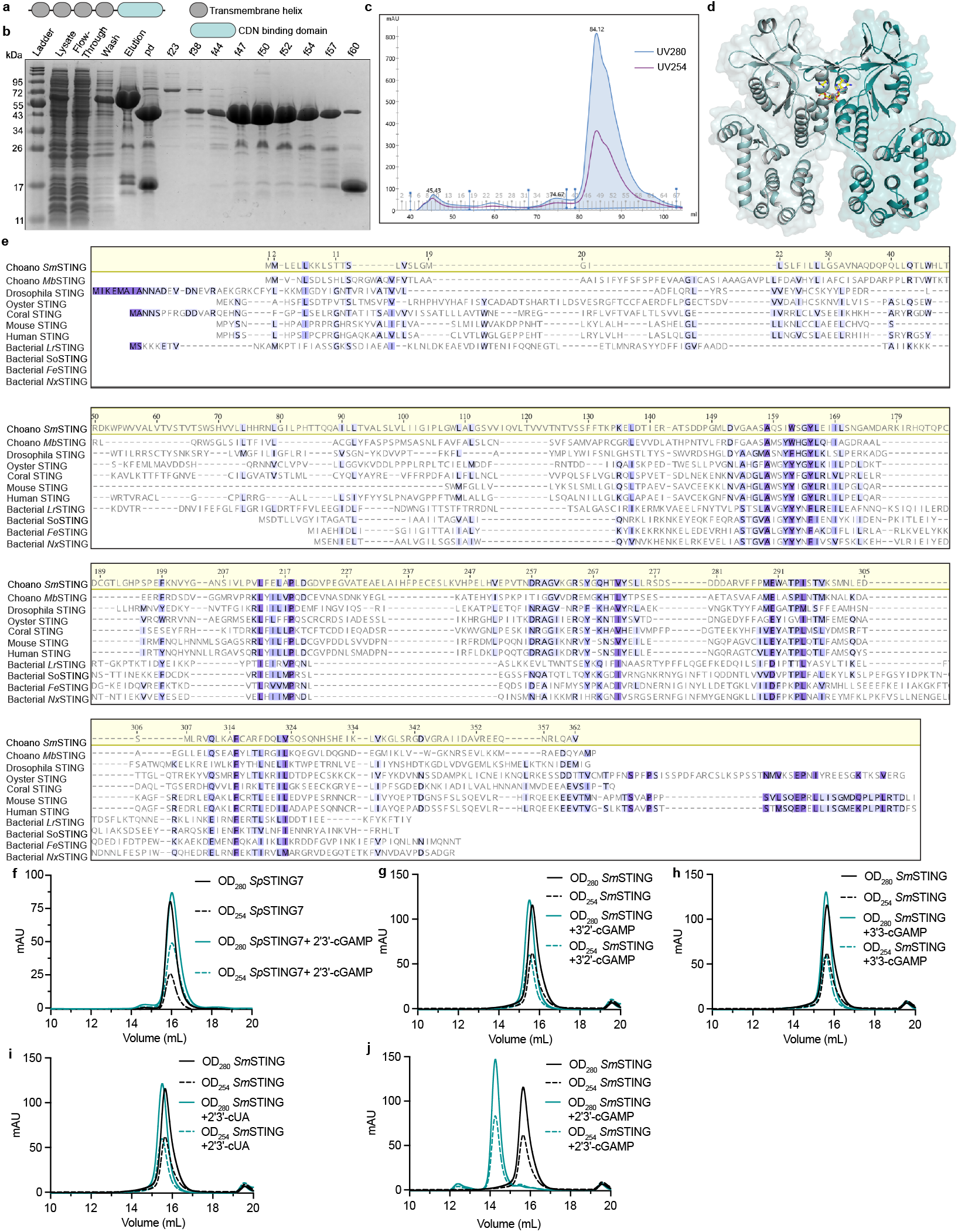
Domain architecture and ligand-induced dimerization of choanoflagellate *Sm*STING. **a**, Domain architecture of *Sm*STING and *Mb*STING, showing four N-terminal transmembrane helices fused to a C-terminal cyclic dinucleotide-binding domain. **b–c**, Purification of *Sm*STING CBD fused to an N-terminal T4 lysozyme for crystallization. **b**, SDS–PAGE with Coomassie stain showing the protein before and after Ni–NTA affinity and size-exclusion chromatography. The elution profile from the final size-exclusion step is shown in **c. d**, Crystal structure of T4 lysozyme–SmSTING CBD in complex with 2′3′-cGAMP at 2.7 Å resolution. **e**, Complete sequence alignment of choanoflagellate, metazoan and bacterial STING receptors. **f**, Size-exclusion chromatograms of coral (*Sp*STING7) CBD in the absence and presence of 2′3′-cGAMP, showing no monomer–dimer transition upon ligand binding. **g–j**, Size-exclusion chromatograms of T4 lysozyme–*Sm*STING CBD in the absence of ligand or in the presence of 3′2′-cGAMP (**g**), 3′3′-cGAMP (**h**), 2′3′-cUA (**i**) or 2′3′-cGAMP (**j**). Dimerization is observed only in the presence of 2′3′-cGAMP.

**Extended Data Figure 4.**
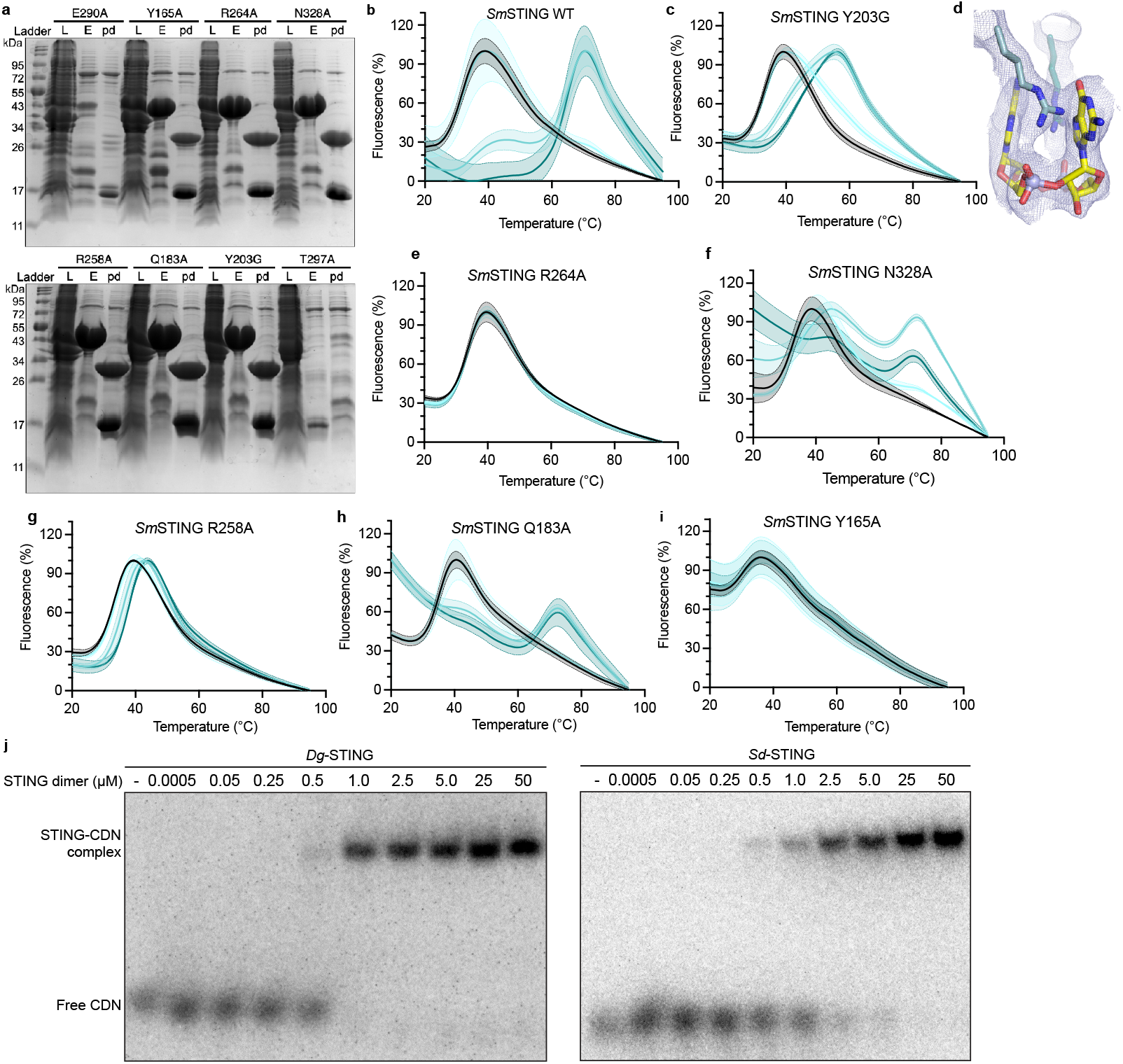
Biochemical characterization of SmSTING and homologs. **a**, Purification of *Sm*STING mutants. Protein expression levels and purity were assessed by SDS–PAGE and Coomassie staining. **b–c**, Normalized fluorescence traces from thermofluor assays, showing changes in fluorescence signal (Y) as temperature (X) increases for wild-type (WT) SmSTING ΔTM and the Y203G mutant. The Y203G mutation disrupts hydrogen bonding between the β-strand lid insertion and the canonical lid. Curves are colored according to ligand concentration, with black indicating no ligand and light to darker teal representing 1, 10, and 100 µM, respectively. Shaded regions surrounding each curve indicate standard deviations (*n* = 3). **d**, Electron density map showing the interaction between residue R264 and the guanine base of 2′3′-cGAMP. **e–i**, Normalized fluorescence traces from thermofluor assays of five additional *Sm*STING mutants in the presence of increasing concentrations of 2′3′-cGAMP. Curves are colored as in panels b–c, and shaded regions indicate standard deviations (n = 3). **j**, Primary EMSA analysis of the binding of bacterial-like choanoflagellate STING receptors *Dg*STING and *Sd*STING to 3′3′-c-di-GMP. Ligand binding was assessed by titrating each STING protein (homodimer form) across concentrations from 0.5 nM to 50 µM (n = 2).

## References

1. Author, A. & Author B. The biology of something: Interesting biology. Biol.Lett. 43, 246–259 (1971)

1. Wein, T. & Sorek, R. Bacterial origins of human cell-autonomous innate immune mechanisms. Nat. Rev. Immunol. 22, 629–638 (2022). 10.1038/s41577-022-00705-4

2. Slavik, K. M. & Kranzusch, P. J. CBASS to cGAS-STING: The Origins and Mechanisms of Nucleotide Second Messenger Immune Signaling. Annu Rev Virol 10, 423–453 (2023). 10.1146/annurev-virology-111821-115636

3. Bernheim, A., Cury, J. & Poirier, E. Z. The immune modules conserved across the tree of life: Towards a definition of ancestral immunity. PLoS Biol. 22, e3002717 (2024). 10.1371/jour-nal.pbio.3002717

4. Aravind, L., Nicastro, G. G., Iyer, L. M. & Burroughs, A. M. The Prokaryotic Roots of Eukaryotic Immune Systems. Annu. Rev. Genet. 58, 365–389 (2024). 10.1146/annurev-genet-111523-102448

5. Ledvina, H. E. & Whiteley, A. T. Conservation and similarity of bacterial and eukaryotic innate immunity. Nat. Rev. Microbiol. 22, 420–434 (2024). 10.1038/s41579-024-01017-1

6. Jenson, J. M. & Chen, Z. J. cGAS goes viral: A conserved immune defense system from bacteria to humans. Mol. Cell 84, 120–130 (2024). 10.1016/j.molcel.2023.12.005

7. Whiteley, A. T. et al. Bacterial cGAS-like enzymes synthesize diverse nucleotide signals. Nature 567, 194–199 (2019). 10.1038/s41586-019-0953-5

8. Cohen, D. et al. Cyclic GMP–AMP signalling protects bacteria against viral infection. Nature 574, 691–695 (2019). 10.1038/s41586-019-1605-5

9. Morehouse, B. R. et al. STING cyclic dinucleotide sensing originated in bacteria. Nature 586, 429–433 (2020). 10.1038/s41586-020-2719-5

10. Lowey, B. et al. CBASS Immunity Uses CARF-Related Effectors to Sense 3′–5′-and 2′–5′-Linked Cyclic Oligonucleotide Signals and Protect Bacteria from Phage Infection. Cell 182, 38–49.e17 (2020). 10.1016/j.cell.2020.05.019

11. Severin, G. B. et al. Activation of a Vibrio cholerae CBASS anti-phage system by quorum sensing and folate depletion. mBio 14, e0087523 (2023). 10.1128/mbio.00875-23

12. Ye, Q. et al. HORMA Domain Proteins and a Trip13-like ATPase Regulate Bacterial cGAS-like Enzymes to Mediate Bacteriophage Immunity. Mol. Cell 77, 709–722.e707 (2020). 10.1016/j.molcel.2019.12.009

13. Jenson, J. M., Li, T., Du, F., Ea, C. K. & Chen, Z. J. Ubiquitin-like conjugation by bacterial cGAS enhances anti-phage defence. Nature 616, 326–331 (2023). 10.1038/s41586-023-05862-7

14. Morehouse, B. R. et al. Cryo-EM structure of an active bacterial TIR– STING filament complex. Nature 608, 803–807 (2022). 10.1038/s41586-022-04999-1

15. Burroughs, A. M. & Aravind, L. Identification of Uncharacterized Components of Prokaryotic Immune Systems and Their Diverse Eukaryotic Reformulations. J. Bacteriol. 202 (2020). 10.1128/JB.00365-20

16. Ko, T. P. et al. Crystal structure and functional implication of bacterial STING. Nat Commun 13, 26 (2022). 10.1038/s41467-021-26583-3

17. Hou, M. H. et al. Structural insights into the regulation, ligand recognition, and oligomerization of bacterial STING. Nat Commun 14, 8519 (2023). 10.1038/s41467-023-44052-x

18. Sun, L., Wu, J., Du, F., Chen, X. & Chen, Z. J. Cyclic GMP-AMP synthase is a cytosolic DNA sensor that activates the type I interferon pathway. Science 339, 786–791 (2013). 10.1126/sci-ence.1232458

19. Burdette, D. L. et al. STING is a direct innate immune sensor of cyclic di-GMP. Nature 478, 515–518 (2011). 10.1038/na-ture10429

20. Ishikawa, H. & Barber, G. N. STING is an endoplasmic reticulum adaptor that facilitates innate immune signalling. Nature 455, 674–678 (2008). 10.1038/nature07317

21. Gao, P. et al. Cyclic [G(2’,5’)pA(3’,5’)p] is the metazoan second messenger produced by DNA-activated cyclic GMP-AMP synthase. Cell 153, 1094–1107 (2013). 10.1016/j.cell.2013.04.046

22. Ablasser, A. et al. cGAS produces a 2’-5’-linked cyclic dinucleotide second messenger that activates STING. Nature 498, 380–384 (2013). 10.1038/nature12306

23. Diner, E. J. et al. The innate immune DNA sensor cGAS produces a noncanonical cyclic dinucleotide that activates human STING. Cell Rep. 3, 1355–1361 (2013). 10.1016/j.celrep.2013.05.009

24. Zhang, X. et al. Cyclic GMP-AMP containing mixed phosphodiester linkages is an endogenous high-affinity ligand for STING. Mol. Cell 51, 226–235 (2013). 10.1016/j.molcel.2013.05.022

25. Ablasser, A. & Chen, Z. J. cGAS in action: Expanding roles in immunity and inflammation. Science 363, eaat8657 (2019). 10.1126/science.aat8657

26. Millman, A., Melamed, S., Amitai, G. & Sorek, R. Diversity and classification of cyclic-oligonucleotide-based anti-phage signalling systems. Nat Microbiol 5, 1608–1615 (2020). 10.1038/s41564-020-0777-y

27. Li, Y. et al. cGLRs are a diverse family of pattern recognition receptors in innate immunity. Cell 186, 3261–3276.e3220 (2023). 10.1016/j.cell.2023.05.038

28. Slavik, K. M. et al. cGAS-like receptors sense RNA and control 3′2′cGAMP signalling in Drosophila. Nature 597, 109–113 (2021). 10.1038/s41586-021-03743-5

29. Holleufer, A. et al. Two cGAS-like receptors induce antiviral immunity in Drosophila. Nature 597, 114–118 (2021). 10.1038/s41586-021-03800-z

30. Culbertson, E. M. & Levin, T. C. Eukaryotic CD-NTase, STING, and viperin proteins evolved via domain shuffling, horizontal transfer, and ancient inheritance from prokaryotes. PLoS Biol. 21, e3002436 (2023). 10.1371/journal.pbio.3002436

31. Woznica, A. et al. STING mediates immune responses in the closest living relatives of animals. eLife 10, e70436 (2021). 10.7554/eLife.70436

32. Richter, D. J., Fozouni, P., Eisen, M. B. & King, N. Gene family innovation, conservation and loss on the animal stem lineage. Elife 7 (2018). 10.7554/eLife.34226

33. Abramson, J. et al. Accurate structure prediction of biomolecular interactions with AlphaFold 3. Nature 630, 493–500 (2024). 10.1038/s41586-024-07487-w

34. Zhou, W. et al. Structure of the Human cGAS–DNA Complex Reveals Enhanced Control of Immune Surveillance. Cell 174, 300–311.e311 (2018). 10.1016/j.cell.2018.06.026

35. Cai, H. et al. The virus-induced cyclic dinucleotide 2′3′-c-di-GMP mediates STING-dependent antiviral immunity in Drosophila. Immunity 56, 1991–2005.e1999 (2023). 10.1016/j.immuni.2023.08.006

36. Kranzusch, P. J. et al. Ancient Origin of cGAS-STING Reveals Mechanism of Universal 2’,3’ cGAMP Signaling. Mol. Cell 59, 891–903 (2015). 10.1016/j.molcel.2015.07.022

37. Carr, M. et al. A six-gene phylogeny provides new insights into choanoflagellate evolution. Mol. Phylogenet. Evol. 107, 166–178 (2017). 10.1016/j.ympev.2016.10.011

38. Jenal, U., Reinders, A. & Lori, C. Cyclic di-GMP: second messenger extraordinaire. Nat. Rev. Microbiol. 15, 271–284 (2017). 10.1038/nrmicro.2016.190

39. Kranzusch, P. J. et al. Structure-guided reprogramming of human cGAS dinucleotide linkage specificity. Cell 158, 1011–1021 (2014). 10.1016/j.cell.2014.07.028

40. Kozlov, A. M., Darriba, D., Flouri, T., Morel, B. & Stamatakis, A. RAxML-NG: a fast, scalable and user-friendly tool for maximum like-lihood phylogenetic inference. Bioinformatics 35, 4453–4455 (2019). 10.1093/bioinformatics/btz305

41. Liu, H. & Naismith, J. H. An efficient one-step site-directed deletion, insertion, single and multiple-site plasmid mutagenesis protocol. BMC Biotechnol. 8, 91 (2008). 10.1186/1472-6750-8-91

42. Vonrhein, C. et al. Data processing and analysis with the autoPROC toolbox. Acta Crystallogr. D Biol. Crystallogr. 67, 293–302 (2011). 10.1107/S0907444911007773

43. Eaglesham, J. B., Pan, Y., Kupper, T. S. & Kranzusch, P. J. Viral and metazoan poxins are cGAMP-specific nucleases that restrict cGAS– STING signalling. Nature 566, 259–263 (2019). 10.1038/s41586-019-0928-6

44. Liebschner, D. et al. Macromolecular structure determination using X-rays, neutrons and electrons: recent developments in Phenix. Acta Crystallographica Section D 75, 861–877 (2019). 10.1107/S2059798319011471

45. Emsley, P. & Cowtan, K. Coot: model-building tools for molecular graphics. Acta Crystallographica Section D 60, 2126–2132 (2004). 10.1107/S0907444904019158

46. Richter, D. J. B. C.; Strassert, J. F. H.; Poh, Y.-P.; Herman, E. K.; Muñoz-Gómez, S. A.; Wideman, J. G.; Burki, F.; de Vargas, C. EukProt: A database of genome-scale predicted proteins across the diversity of eukaryotes. Peer Community Journal 2, article no. e56 (2022). 10.24072/pcjournal.173

47. Priyam, A. et al. Sequenceserver: A Modern Graphical User Interface for Custom BLAST Databases. Mol. Biol. Evol. 36, 2922–2924 (2019). 10.1093/molbev/msz185

48. Le, S. Q. & Gascuel, O. An improved general amino acid replacement matrix. Mol. Biol. Evol. 25, 1307–1320 (2008). 10.1093/molbev/msn067

49. Letunic, I. & Bork, P. Interactive Tree of Life (iTOL) v6: recent updates to the phylogenetic tree display and annotation tool. Nucleic Acids Res. 52, W78–W82 (2024). 10.1093/nar/gkae268

50. Krogh, A., Larsson, B., von Heijne, G. & Sonnhammer, E. L. Predicting transmembrane protein topology with a hidden Markov model: application to complete genomes. J. Mol. Biol. 305, 567–580 (2001). 10.1006/jmbi.2000.4315

