## Supplementary Table 2 for "A choanoflagellate cGLR-STING pathway reveals evolutionary links between bacterial and animal immunity"

**Supplementary Table 2. Crystallographic Statistics, related to Fig. 3**

| | *Sm*STING–2′3′-cGAMP (SeMet) | *Sm*STING–2′3′-cGAMP |
|---|---|---|
| **Data Collection** | | |
| Resolution (Å)[a] | 75.90–3.32 (3.50–3.32) | 34.04–2.65 (2.71–2.65) |
| Wavelength (Å) | 0.9792 | 0.92010 |
| Space group | P 1 $2_1$ 2 | P 1 $2_1$ 2 |
| Unit cell: a, b, c (Å) | 66.39, 88.30, 76.15 | 66.95, 88.74, 76.02 |
| Unit cell: α, β, γ (°) | 90.0, 94.6, 90.0 | 90.0, 94.1, 90.0 |
| Molecules per ASU | 2 | 2 |
| Total reflections | 82710 (8528) | 184837 (13747) |
| Unique reflections | 12067 (1379) | 25932 (1855) |
| Completeness (%)[a] | 92.2 (72.9) | 99.7 (96.8) |
| Multiplicity[a] | 6.9 (6.2) | 7.1 (7.4) |
| $I/\sigma I$[a] | 5.4 (0.6) | 4.51 (0.77) |
| CC(1/2)[b] (%)[a] | 99.2 (38.4) | 98.4 (37.8) |
| Rpim[c] (%)[a] | 11.2 (155.2) | 11.3 (84.6) |
| Sites | 20 | |
| **Refinement** | | |
| Resolution (Å) | | 34.04–2.65 |
| Free reflections | | 2024 (145) |
| R-factor / R-free | | 24.3 / 27.5 |
| Bond distance (RMS Å) | | 0.003 |
| Bond angles (RMS °) | | 0.63 |
| **Structure/Stereochemistry** | | |
| No. atoms: protein | | 5759 |
| No. atoms: ligand | | 45 |
| No. atoms: solvent | | 79 |
| Average B-factor: protein | | 71.96 |
| Average B-factor: ligand | | 60.80 |
| Average B-factor: water | | 56.81 |
| Ramachandran plot: favored | | 98.04% |
| Ramachandran plot: allowed | | 1.96% |
| Ramachandran plot: outliers | | 0.00% |
| Rotamer outliers | | 0.65% |
| MolProbity[d] score | | 1.46 |
| Protein Data Bank ID | | 9Q1F |

[a] Highest resolution shell values in parenthesis
[b] (Karplus and Diederichs, 2012)
[c] (Weiss, 2001)
[d] (Chen et al., 2010)
